# A Primary Cell Culture Platform for Studying Venom Gland and Brain Tissue in Octopus

**DOI:** 10.64898/2026.06.01.729309

**Authors:** James Vincent Parziale, Saurabh Attarde, Fatima Khalid, Pollena Devi Sangana, Mandë Holford

**Author notes:** Corresponding author: Mandë Holford. These authors contributed equally (James Vincent Parziale, Saurabh Attarde).

## Abstract

Coleoid cephalopods, squids, cuttlefish, and octopuses, have emerged as powerful model organisms for studying neurobiology, development, and behavior, however, cellular tools for investigating their specialized tissues remain limited. In particular, their venom producing gland, the posterior salivary gland (PSG), has been extensively described anatomically and histologically, yet remains largely inaccessible to experimental investigation at the cellular level. Here, we report the first establishment of primary cell cultures derived from both the optical lobe and PSG tissues of *Octopus bimaculoides*. Building on recent advances in cephalopod brain cultures, we adapted and optimized dissociation and culture conditions to support short-term survival and attachment of cells *in vitro*. We show that passive cell release during tissue handling, rather than enzymatic treatment, yields viable cultures from both tissues, and poly-D-lysine markedly improves the adherence of PSG-derived cells. Morphological analyses and fluorescent staining confirm the presence and viability of distinct cell populations, while cell cycle analysis indicates that the majority of cells reside in G_0_/G_1_ phase. Notably, *O. bimaculoides* brain cultures exhibit features comparable to those previously described in squid, suggesting conserved aspects of coleoid cellular physiology. Together, our findings establish a foundational *in vitro* platform for studying octopus PSG and neural cell biology, providing a tractable system for probing venom biosynthesis, secretion, and neural regulation in coleoid cephalopods.

## Introduction

Since the early 20^th^ century, coleoid cephalopods have been an exciting model to investigate neuroscience, behavior, and developmental biology.^1,2^ Coleoid cephalopods, or soft-bodied cephalopods, include octopuses, squids, and cuttlefish. Coleoid’s rapid development, well-defined embryonic stages, and tractable laboratory husbandry have allowed species such as *Octopus bimaculoides* (California two-spot octopus), *Sepia officinalis* (common cuttlefish), and *Euprymna berryi* (hummingbird bobtail squid) to play instrumental roles as emerging research model organisms. In recent years, the establishment of genomic and transcriptomic resources, the emergence of CRISPR-based gene editing, and successful transgenesis in coleoid embryos have further positioned cephalopods as a robust system for investigating developmental and neural mechanisms.^3^

Unlike shelled mollusks, like nautiluses, coleoids have either an internalized shell known as the cuttlebone (in cuttlefish), a reduced chitinous pen (squids), or full loss of their shell (octopus). This morphological change is accompanied by high degree of specialization in their soft tissues aiding in survival such as camouflage, flexible musculature allowing for intricate movement manipulation with their limbs, and predatory efficiency through mechanical means and through the aid of venom.^4–6^

The diversity of cephalopod venoms is understudied, limiting our understanding of this class of compounds.^5,6^ Generally, coleoid cephalopod venom compounds, referred to as cephalotoxins, are found in their posterior salivary gland (PSG) and are chemically heterogeneous, composed of larger and often glycosylated proteinaceous factors, with different enzymatic activities.^4–6^ The few cephalotoxins that have been identified are a mix of biologically active compounds that predominantly act as neurotoxins, with a few others exerting effects related to hypotension or antibacterial activity.^9,10^ There is only one known structural classification of cephalotoxins from *Sepia esculenta*, SE-CTX. SE-CTX has a 1052-amino-acid sequence and contains EGF-like, Sushi, TSP type-1, and LDL-receptor class A domains.^9, 10^ Because the EGF-like domain is the only domain commonly found in other venoms, particularly in cnidarians, it is currently classified as an EGF-neurotoxin, but its mechanism of action is unknown. In addition to cephalotoxins, tetrodotoxin (TTX), a small non-peptidic sodium-channel blocker, is found in the PSG of blue-ringed octopuses (*Hapalochlaena* spp.) and many other marine organisms. TTX, which is fatal, is synthesized by endosymbiotic bacteria and sequestered and stored in concentrated amounts in the PSG.^11,12^

The venom-producing PSG is dorsal to the mantle cavity behind the cranium and connects to the buccal mass via the posterior salivary gland duct (**Figure 1A)**. Cephalopod buccal mass is composed of a chitinous beak and radula, and muscle tissue. During predation cephalopods use their beak to drill and bite into their target’s tissue and introduce saliva that includes a mixture of venomous secretions and digestive enzymes released from their PSG. There is anatomical variation among octopus, squid, and cuttlefish PSGs. The cuttlefish and octopus, such as *O. bimaculoides,* have pairs of PSGs, whereas most squids have one PSG (**Figure 1A)**.^6–8^ Previous work has characterized the PSG as a branched tubular organ supported by a framework of connective and vascular tissue. The precise mode of secretion within the PSG has yet to be fully elucidated, though current evidence suggests a merocrine or apocrine mechanism.^6,8^

**Figure 1:**
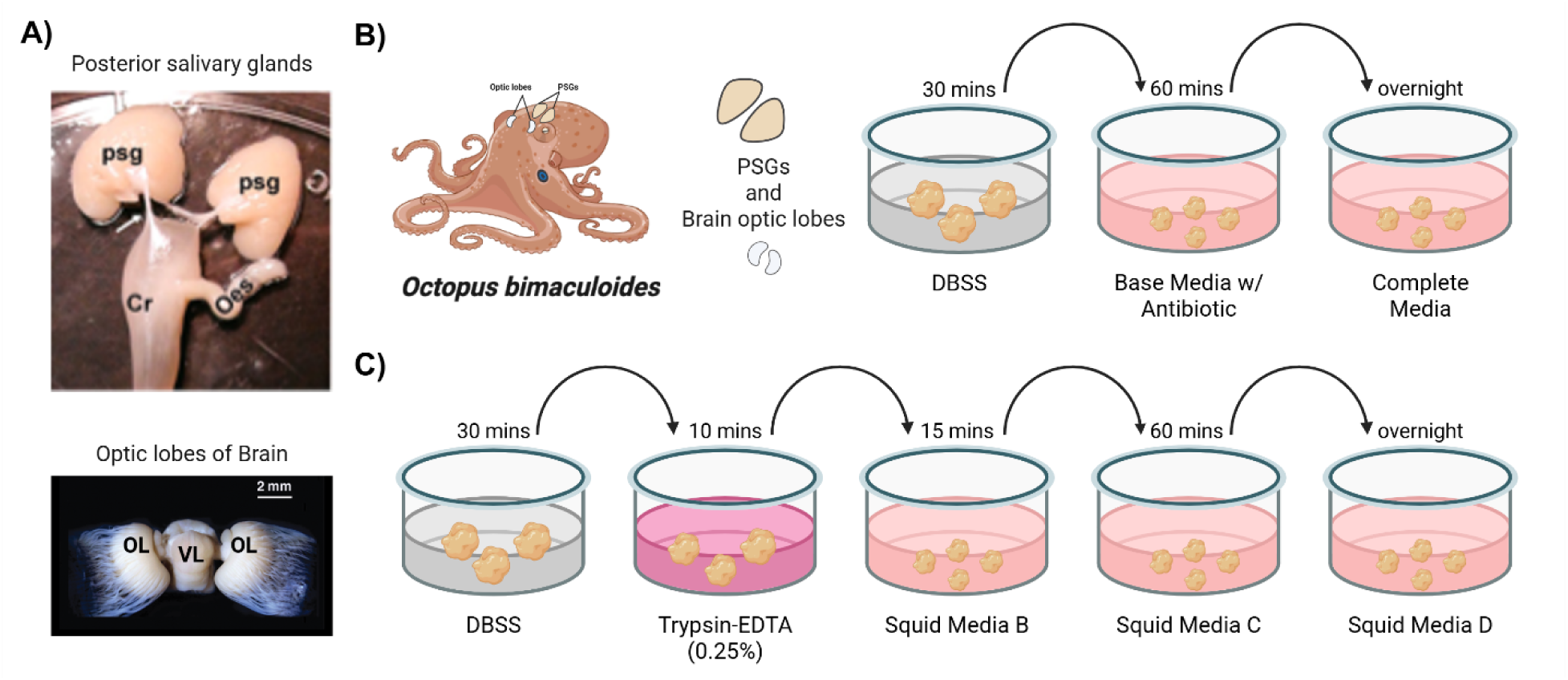
Experimental strategy for generation of cephalopod *in vitro* cell models. A) Posterior salivary glands (PSGs) and optic lobes dissected from *Octopus bimaculoides* (PSG image adapted from Baldascino *et al*., 2017^16^; Brain image adopted from Messenger and Hanlon, 2018^17^); psg: posterior salivary gland, Cr: crop, Oes: oesophagus, OL: optic lobe, and VL: vertical lobe. B) Optimized *O. bimaculoides* tissue processing workflow. Dissected PSGs and brain optic lobes are transferred sequentially in a 24-well plate using sterile tweezers: first into disinfection balanced salt solution (DBSS) for 30 minutes, followed by base media supplemented with antibiotic for 60 minutes, and finally into complete media for overnight incubation (See Methods for formulation of media types). C) Original tissue dissociation workflow adapted from Kim et al 2025^15^. Tissue is incubated in DBSS (30 minutes), followed by 0.25% Trypsin-EDTA (10 minutes), then transferred sequentially into Squid Media B (15 minutes), Squid Media C (60 minutes), and finally Squid Media D for overnight culture. Cells dissociate progressively during these steps without additional mechanical disruption. For both protocols, on Day 1, cells are maintained in Complete media/Squid Media D and residual tissue is removed from the final well.

The optic lobe is an important neural structure in cephalopods. It makes up a large part of the brain and is involved in processing visual information and integrating sensory and motor information.^13^ It has separate cortical and medullary areas, and its layered cellular structure reportedly supports complex visual behaviors and learning.^13^ The cortical region contains densely packed small interneurons arranged in layered circuits for early visual processing, while the medulla comprises larger projection and integrative neurons that link visual input to motor outputs. ^13^ (**Figure 1A)**.

While previous reports have characterized morphological and molecular components of cephalopod PSG venom glands and brain optic lobes, there is a dearth of *in vitro* approaches for investigating cephalopod cell biology. Most functional studies rely on whole embryos or intact adult tissues, which are not readily applicable for systemically elucidating cellular mechanisms underlying neural differentiation, gland development, or specialized secretory functions. As in other marine invertebrate phyla, long-term primary culture and continuous cell line generation have been technically challenging due to difficulties in optimizing osmolarity, extracellular matrix requirements, growth factor dependencies, and axenic culture conditions.^14^ Consequently, the field lacks scalable model systems for genetic manipulation, controlled perturbation of signaling pathways, or high-throughput functional assays of cephalopod-specific genes including many that remain unannotated. A recent report described the successful establishment of primary brain tissue cultures from the coleoid cephalopod *Euprymna berryi*^15^. Cephalopod brain tissue differs in cellular composition and function from venom gland tissues; however, as *E. berryi* is a venomous molluscan species, it is possible to share methodological parallels, including similar strategies for tissue dissociation and reliance on comparable growth and culture parameters for success.

Here we present the first octopus optical lobe and PSG primary cell cultures, which can be used to advance our understanding of octopus brain and venom systems. We selected *Octopus bimaculoides* as the research organism for this study due to its accessibility and the large size of its optical lobes and venom PSGs. Our established optic lobe and PSG primary cell culture systems from *O. bimaculoides*, serve as a foundational platform for future cellular and molecular investigations of cephalopod neuronal and venom gland biology, which remains largely understudied. Interestingly, our identified primary brain cultures from *O. bimaculoides* closely resemble the previously established *Euprymna berryi* brain cultures from Kim *et al*. (2025),^15^ highlighting potential parallels in cephalopod cellular physiology that could inform future studies in cephalopod development and neurobiology.

## Results

To obtain *O. bimaculoides* primary cultures, we modified the approach and media formulations described by Kim et al. (2025)^15^ for optical lobe brain tissue of squid *E. berryi* (**Figure 1B**). Specifically, we omitted the trypsin-EDTA dissociation step and instead used sequential incubations in base and complete media following DBSS washes to achieve tissue processing without enzymatic digestion. We applied our optimized protocol to both optical lobe brain (**Figure 2A**) and venom PSG (**Figure 2B**) tissue cultures. Growth conditions were monitored over 5 days (Figure 2). Over this time, we observed cells with clearly defined nuclei and discernible cellular morphologies, including spherical and fibroblast-like structures in the optical lobe brain tissue (**Figure 2A**), while the venom PSG tissue yielded only spherical cells (**Figure 2B**).

**Figure 2:**
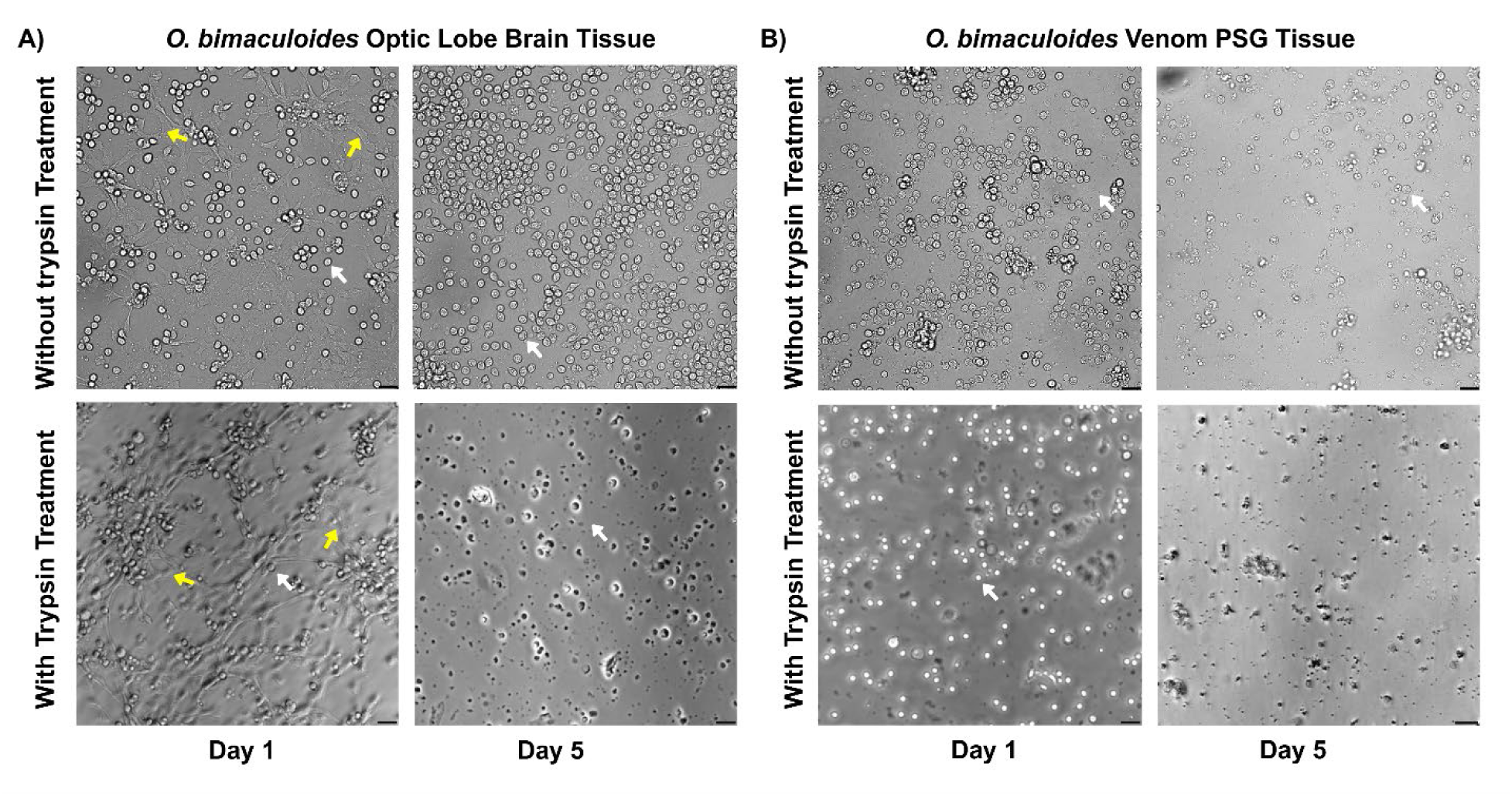
Dissociation of *O. bimaculoides* optical lobe brain and venom PSG tissue does not require trypsin. Optical lobe brain and venom PSG tissue from *O. bimaculoides* was dissociated without and with trypsin to determine optimal conditions for dissociation. Tissues were imaged on days 1-5 after plating. A) Optical lobe brain cells dissociated without trypsin treatment exhibit more fibroblast-like cells (yellow arrows) on day 1. By day 5 only spherical cells (white arrows) are observed. Tissue dissociated with trypsin appears to yield both spherical cells and fibroblast-like cells from day 1 to day 5. Scale bar = 50 µm. B) Venom PSG tissue was also dissociated with and without trypsin and monitored from day 1 to day 5. Unlike optical lobe brain cells which included a mixture of spherical and fibroblast-like cells, PSG tissue yielded predominantly spherical cells. Similar to optical lobe brain cells dissociation of venom PSG tissue without trypsin appear to improve the cell density. Images were captured with a 20x objective. Scale bar = 50 µm.

To further characterize temporal changes in cell behavior independent of trypsin treatment, cultures generated without trypsin were monitored from Day 1 to Day 5 under DBSS, base media, and complete media conditions (**Supplementary Figures 1 and 2**). On Day 1, a high proportion of Hoechst-positive nuclei was observed across all conditions, indicating strong initial viability (**Supplementary Figures 1 and 2**). Optical lobe brain cells were moderately dense and largely well-dispersed, with minimal aggregation. Morphologically, >80% of optical lobe cells appeared spherical, while a smaller fraction (<20%) of elongated, fibroblast-like cells was observed in brain-derived cultures (**Figure 2A**). Similarly, venom PSG-derived cultures from *O. bimaculoides* on Day 1 exhibited moderate cell density with cells primarily well-dispersed across the field of view. The majority (>95%) of PSG cells displayed a spherical morphology with clearly defined boundaries, and little to no presence of elongated or fibroblast-like cells was observed (**Figure 2B**).

By Day 2, apparent cell density increased across all conditions, most prominently in DBSS, where clustering of optical lobe brain and venom PSG spherical cells resulted in localized high-density regions (**Supplementary Figures 1 and 2**). In optical lobe brain cultures, the proportion of fibroblast-like cells increased relative to Day 1, becoming more readily detectable across multiple experimental replicates (**Figure 2A; Supplementary Figure 1**), whereas venom PSG cultures remained predominantly (>90%) spherical (**Figure 2B; Supplementary Figure 2**). By Day 3, DBSS conditions exhibited the highest apparent cell density for optic lobe brain, largely due to continued aggregation rather than an increase in total cell number (**Supplementary Figures 1**). In contrast, base and complete media showed a relative reduction in cell density and increased dispersion.

From Day 4 onward, a progressive decline in cell density and viability was observed across all media conditions for both optical lobe brain and venom PSG tissues, as evidenced by a reduction in Hoechst-positive nuclei per field and increased debris-like structures indicate cell death (**Supplementary Figures 1 and 2**). By Day 5, total cell numbers were visibly reduced for both optic lobe brain and venom PSG, particularly in base and complete media, where sparse and dispersed cells predominated (**Supplementary Figures 1 and 2**). DBSS cultures appear to maintain comparatively higher apparent density; however, this may be due to aggregation of cells rather than sustained proliferation. In brain optical lobe cultures, fibroblast-like cells persisted but appeared reduced in number, likely due to detachment over time (**Figure 2A**), while venom PSG cultures remained largely spherical with diminished structural integrity (**Figure 2B**).

As shown in Figure 2A, brain optical lobe cultures exhibited two morphologically distinct cell types fibroblast-like and roughly spherical cells, in both trypsin-treated and non-trypsin-treated conditions. A higher relative abundance of fibroblast-like cells was observed in trypsin-treated wells on Day 1; however, by Day 5, the overall morphology and relative proportions of cell types appeared similar between conditions. In contrast, venom PSG cultures (**Figure 2B**) displayed a single predominant spherical morphology, with non-trypsin-treated conditions showing higher apparent cell density and improved preservation of cell structure. Comparison on Day 5 of optical lobe brain and venom PSG tissues emphasize the differences in cell morphologies, suggesting their different functions. Visual estimation of cell counts suggest that dissociation of optical lobe brain and venom PSG tissue without trypsin increased the number of cells present (149 for brain cells with trypsin vs 856 for brain cells without trypsin, and 98 for PSG cells with trypsin vs 168 for PSG cells without trypsin) (**Figure 2A-2B**). The tendency of cells to aggregate and detach in primary cultures limited traditional quantification of standard cell viability assays, therefore the cell counts presented are estimated. Figure 3 provides higher-magnification visualization confirming nuclear integrity and cellular morphology, including the presence of fibroblast-like cells in optical lobe brain cultures (**Figure 3A**) and predominantly spherical cells in venom PSG cultures (**Figure 3B**).

**Figure 3:**
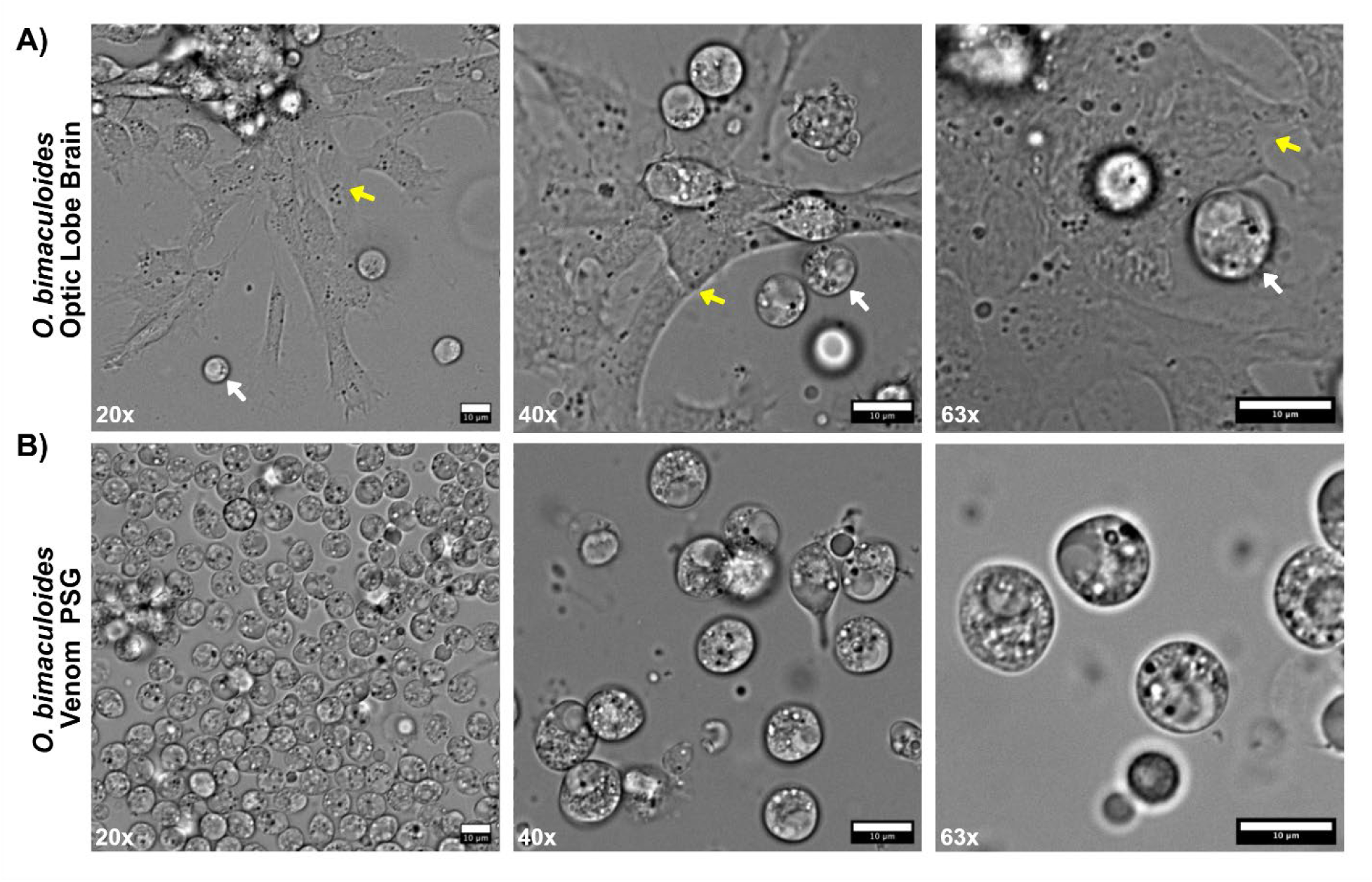
Dissociated *O. bimaculoides* brain and PSG tissue display distinct cell types. Optical lobe brain and PSG tissues were dissociated without trypsin and imaged on day 1 after plating at 20x (left). 40x (middle) and 63x (right). A) Brain cells consist of fibroblast-like cells (yellow arrows) and roughly spherical cells (white arrows). B) gland cells are roughly spherical cells. Scale bar = 10 µm.

Despite successful dissociation, cell attachment remained low across both tissue types, with relatively few adherent cells observed per field (**Figure 4**). Attachment improved in optical lobe brain cultures in regions enriched with fibroblast-like cells (**Figure 4A**). The addition of poly-D-lysine (PDL) significantly increased the proportion of adherent spherical cells in both optical lobe brain and venom PSG cultures (**Figure 4A-4B**). Specifically, with base media, approximately 912 cells were observed for optical lobe brain cultures with PDL compared to 321 without PDL, and 742 cells for venom PSG cultures with PDL compared to 186 without PDL (**Figure 4A-4B**). Addition of PDL appears to promote cell spreading and polarity compared to predominantly rounded, loosely associated cells observed under non-coated conditions. Fibroblast-like cells, however, appeared to exhibit reduced attachment on PDL-coated surfaces (**Figure 4A**).

**Figure 4:**
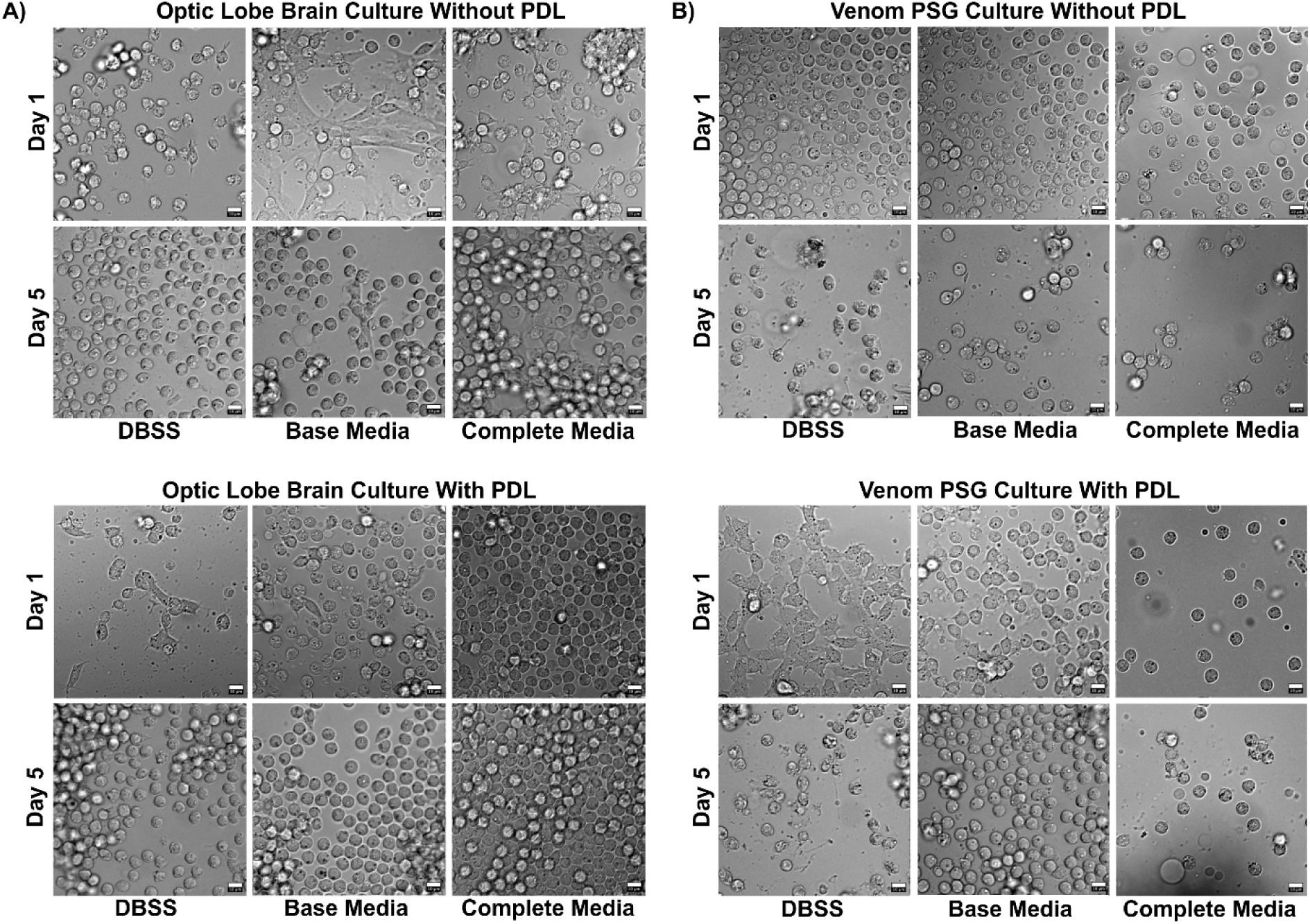
Dissociated optical lobe brain and venom posterior salivary gland (PSG) cells show improved attachment on PDL-coated plates. (A) Optical lobe brain and (B) Venom PSG tissues were dissociated and imaged on Day 1-5 on culture plates that were either uncoated or coated with 50 µg/mL poly-D-lysine (PDL). Dissociated optical lobe brain cells were roughly spherical and showed improved attachment on PDL-coated plates on day 1, although PDL had an adverse effect on optical lobe brain fibroblast-like cells. Over time, optical lobe brain cells detached and aggregated under both PDL and non-PDL conditions. Dissociated venom PSG cells also exhibited improved attachment on PDL-coated plates on Day 1, but cells detached from PDL-treated plates by Day 5, with cell density decreasing over time in both treatment conditions. Venom PSG cultures exhibited the highest cell density in DBSS and base media with antibiotics, the lowest density in complete media, and overall best attachment in DBSS. Images were captured with a 20x objective. Scale bar = 10 µm.

To better assess cell structure, confirm species identity, and evaluate compatibility with additional staining protocols, we performed cphalloidin and Hoechst staining. In optical lobe brain cultures, we observe two distinguishable morphologies: roughly spherical cells with slight polarity and fibroblast-like cells confirming what was previously identified in Figures 2 and 3 brightfield microscopy **(Figure 5A)**. In venom PSG-derived cultures, only one clear attached and polarized cell-type was identified, similarly confirming what was previously identified in Figures 2 and 3 brightfield microscopy **(Figure 5A)**. The average cell diameter of the roughly round optical lobe brain cells was 9.60 µm +/- 0.62 µm. The average cell diameter of the roughly round venom PSG cells was 9.37 µm +/- 0.67 µm.

**Figure 5:**
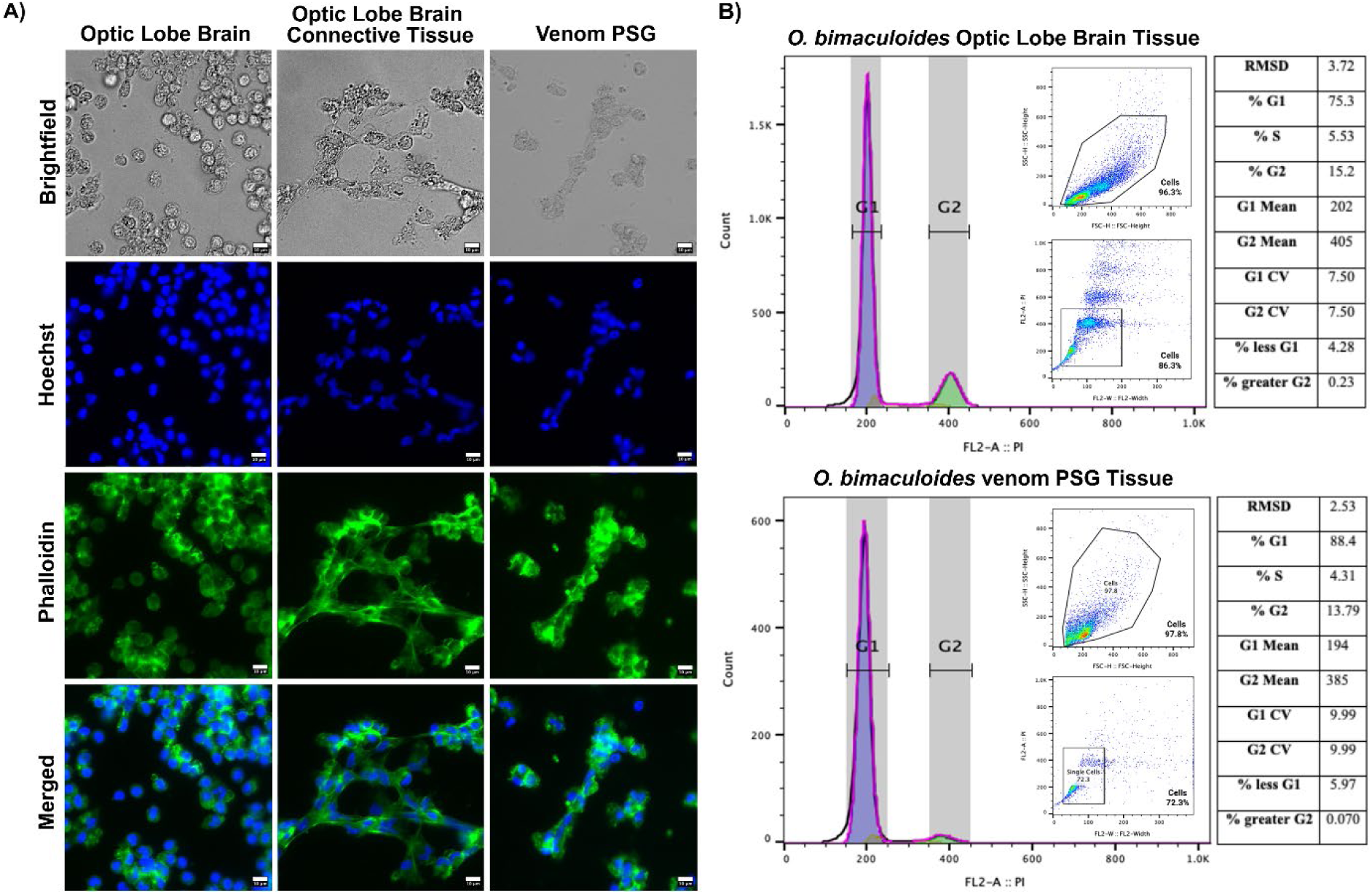
DNA and F-actin staining identify tissue-specific cell types in dissociated *O. bimaculoides* optical brain and gland tissue, with the majority of cells residing in the G_0_/G_1_ states. **A)** From left to right brightfield, Hoechst, phalloidin and merge the Hoecht and phalloidin staining of dissociated and cultured brain, brain connective tissue, and gland tissue. Hoechst staining in blue reveals nuclei and phalloidin in green reveals actin filaments Images captured with a 20x objective. Scale bar = 10 µm. **B)** Flow cytometry cell cycle analysis of optical lobe brain cells (top) venom cells (bottom) stained with propidium iodide (PI). Histograms show cell counts versus PI fluorescence intensity, corresponding to DNA content. Cell cycle distribution (G0/G1, S, and G2 phases) was modeled using the Watson pragmatic model. Insets display gating strategies for singlet cell populations. Tables summarize the percentage of cells in each phase, along with root mean square deviation (RMSD) and coefficient of variation (CV) values indicating goodness of fit. A total of 20,000 events were analyzed for brain tissue and 10,000 events for PSG tissue, with the majority of cells residing in the G0/G1 phase.

To determine whether cells were viable and/or dividing in culture, we performed flow cytometry, **(Figure 5B)**. Using propidium iodide (PI) staining for cell cycle analysis, we found that optical lobe brain cultures contained a higher proportion of cells in the G_0_/G_1_ phase (G_0_/G_1_ = ∼79.58, S = ∼5.53%, G_2_/M = ∼15.43%). The venom PSG samples showed overall lower cell counts, but exhibited a similar distribution, with most cells also in G_0_/G_1_ (G_0_/G_1_ = ∼94.37%, S = 4.31%, G_2_/M = ∼3.72). Slight deviance from 100% totals on the cell counts may be due to aneuploidy, aggregates, or mixed cell populations.

## Discussion

Cell-based model systems in mollusks are challenging to develop and have difficulties such as obtaining proliferative vs quiescent axenic cells, osmolarity and chemical composition of growth media, optimal adhesion, dissociation, and population^18^. Until the recent husbandry breakthrough by the Marine Biological Laboratory to culture various species of cephalopods, laboratory recreation of marine mollusk breeding conditions were challenging, expensive, and time-consuming^19^. In scientific venues, venoms have transformed from a deadly force of nature to a powerful ally in medicine and biotechnology^20^. Venoms are increasingly recognized as the foundation for a new generation of life-saving therapeutics, chemical probes, and even pesticides^21,22^. Venomous mollusks, particularly cone snail and cephalopods, possess complex venom compositions and secretion systems that remain largely uncharacterized at the cellular and molecular levels^6,23,24^. While a variety of computational and analytical techniques can generate predictive information about molluscan venom components, there is still a lack of robust, and reliable experimental cell models for studying these complex systems. This absence has contributed to persistent knowledge gaps regarding how these venoms evolve, develop and function across molluscan lineages. Additionally, for venomous gastropods, their shells present a physical barrier to experimental manipulation, which is why we intentionally focus on cephalopods as an emerging venom research model.

Our study represents the first established primary cell cultures of *O. bimaculoides* venom posterior salivary gland (PSG) and brain optical lobe tissues. The need for such models is compelling, as there are no existing cellular approaches that allow for functional and molecular studies of venom that are experimentally tractable, cost-effective, promote conservation of animals, and are reproducible. *O. bimaculoides* are a molluscan lineage whose venom systems remain experimentally inaccessible despite strong molecular and cellular interest.

Octopus and other coleoid cephalopods contain soft venom PSG and optical lobe brain tissue, whose cells are readily dissociated without enzymatic manipulation (**Figures 2 & 3**). Furthermore, *O. bimaculoides* brain cultures demonstrated substrate-specific preferences: spherical cells adhered more strongly to PDL-coated plates, while fibroblast-like cells were more apparent on non-PDL surfaces (**Figure 4**). Consistent with other molluscan systems, dissociation strategy and cell adhesion substrate were critical determinants of culture success and influenced which cell types survived.^14,25,26^ Our findings suggest that, depending on tissue type and the populations of interest, extensive enzymatic dissociation with trypsin may be unnecessary, or too harsh for establishing healthier cephalopod cultures. Although the cell cultures survived only for 5 days and were not proliferative, they all maintained membrane bound organelles with an identifiable nucleus consistent with viable cells (**Figures 4 & 5, Supplementary Figure 3**). This suggests that *O. bimaculoides* venom PSG and optical lobe brain tissue may be amenable to long term *in vitro* cell culture models with optimization.

While it was possible to improve dissociation efficiency for *O. bimaculoides*, cell attachment remains a major bottleneck. Additional limitations of this study include issues common to other primary cell cultures, namely variations in cell distribution and attachment, which resulted in a number of qualitative evaluations. Extended analyses were also limited by the culture’s short lifespan of only five days. Identifying the precise extracellular matrix components that octopus venom gland epithelial cells rely on may help recreate more physiologically relevant environments. Future work should test additional substrates such as laminin, collagen, fibronectin, and three-dimensional matrices such as Matrigel, any of which may support distinct cell subtypes. Improving staining and characterization protocols is also essential, as conventional buffers and hybridization conditions may damage molluscan cells or alter probe binding due to differences in osmolality or membrane composition. Additionally, there is a need to address the low yield of actively dividing cells for both optical lobe brain and venom PSG tissues.

Mammalian primary and immortalized cell models have steadily improved because researchers draw from extensive genomic, transcriptomic, proteomic, and metabolomic datasets^27^. Similarly, integrating existing experimental cell culture techniques with genomics, transcriptomics, proteomics, histology, and other venomics approaches will be important for making advances to culture venomous organisms^28,29^. As marine molluscan cell culture methods continue to advance, we will gain a deeper understanding of the unique biology of venomous marine mollusks and their specialized secretory tissues. Ultimately, improved culture systems will enable detailed visualization and manipulation of the molecular pathways underlying venom biosynthesis, packaging, and release, opening avenues for novel discovery, and for functional studies for molluscan venom models that have long remained out of reach.

## Methods

The tables below outline the different media used to generate *O. bimaculoides* cell cultures.

### *O. bimaculoides* PSG Dissection

Wild-caught *O. bimaculoides* were provided by the Marine Biological Laboratory at Woods Hole, Massachusetts. A total of 45 adult animals of both sexes were used in this study. Animals were maintained under standard husbandry conditions at the Marine Biological Laboratory, including recirculating seawater systems with controlled temperature (approximately 16–18 °C), salinity (∼32–35 ppt), and a natural or simulated light–dark cycle. Animals were housed individually and monitored regularly prior to experimentation. Prior to tissue collection, animals were deeply anesthetized by immersion in a 1:1 dilution of 2-4% ethanol prepared in home tank seawater. Depth of anesthesia was confirmed by the complete loss of righting reflex and the absence of responses to tactile stimulation. Following confirmation of deep anesthesia, animals were humanely euthanized by decerebration (brain destruction) prior to tissue dissection, in accordance with established cephalopod guidelines. Dissected tissues were used for primary culture and downstream analyses, with experiments performed across multiple biological replicates and technical replicates where applicable (including three biological and three technical replicates for flow cytometry experiments). No formal randomization or blinding procedures were applied, as the study was exploratory in nature and focused on methodological development. All experiments involving live cephalopods were reviewed and approved by the Institutional Animal Care and Use Committee (IACUC) at the Marine Biological Laboratory (MBL) and were conducted in strict accordance with the MBL Cephalopod Care and Use Policy.

### *O. bimaculoides* Brain and Gland Cell Dissociation and Culture

A modified version of the protocol in Kim *et al*. (2025)^15^ to culture the optical lobe tissue of *Euprymna berryi* was used to culture both brain optic lobe and venom PSG cells of *O. bimaculoides* over 9 experiments **(Figure 1B**). The tissue plated directly in DBSS (1 mL) was allowed to sit for 30 minutes. No enzymatic or active mechanical dissociation (e.g., trituration or mincing) was performed. Instead, cells were released passively over time during sequential transfers between solutions. Each piece of tissue (entire brain optic lobe and venom PSG) was moved with sterile forceps to a well with base media with antibiotic (1 mL) **(Table 2)** for 1 hour. The tissue was moved to a new well with complete media (1 mL) **(Table 3)** overnight at 18°C. All culture media were maintained at a pH of approximately 7.6–7.8 and an osmolarity of ∼950–1050 mOsm/kg, consistent with seawater conditions for marine invertebrates. The next day the tissue was removed, and new media was replenished in the complete media well and cultures were monitored over time. Brightfield imaging was performed daily across multiple fields of view within each well to monitor cell behavior and morphology. Media change occurred when phenol red indicator turned yellow on day 4. Brightfield images were taken on a Zeiss Axio Observer 7 and processed on FIJI (V.2.9.0/1.53t).^30^ Cell sizes of around 200 cells were measured in Fiji using the straight-line tool using manual measurement of major and minor axes, measured as area and converted to equivalent diameter.

**Table 1:** Disinfection Balanced Salt Solution (1 Liter) ^15^

| **Table 1: Disinfection Balanced Salt Solution (1 Liter)<sup>15</sup>** |  |  |
| --- | --- | --- |
| NaCl | 26.0 g | 445 mM |
| KCl | 1.08 g | 14.5 mM |
| MgCl<sub>2</sub> • 6H<sub>2</sub>O | 4.70 g | 23.1 mM |
| MgSO<sub>4</sub> • 7H<sub>2</sub>O | 6.52 g | 26.5 mM |
| NaHCO<sub>3</sub> | 0.30 g | 3.57 mM |
| NaH<sub>2</sub>PO<sub>4</sub> • 2H<sub>2</sub>O | 0.06 g | 0.385 mM |
| CaCl<sub>2</sub> • 2H<sub>2</sub>O | 1.48 g | 10.1 mM |
| Glucose | 0.3 g | 1.67 mM |
| Distilled H<sub>2</sub>O | Dissolve above in 1L | - |
| Penicillin-Streptomycin<br>(10,000 U/mL) (Cytiva, USA) | Add to smaller aliquot when<br>ready to use | Add 0.5% (v/v) and filter<br>prior to use |

**Table 2:**
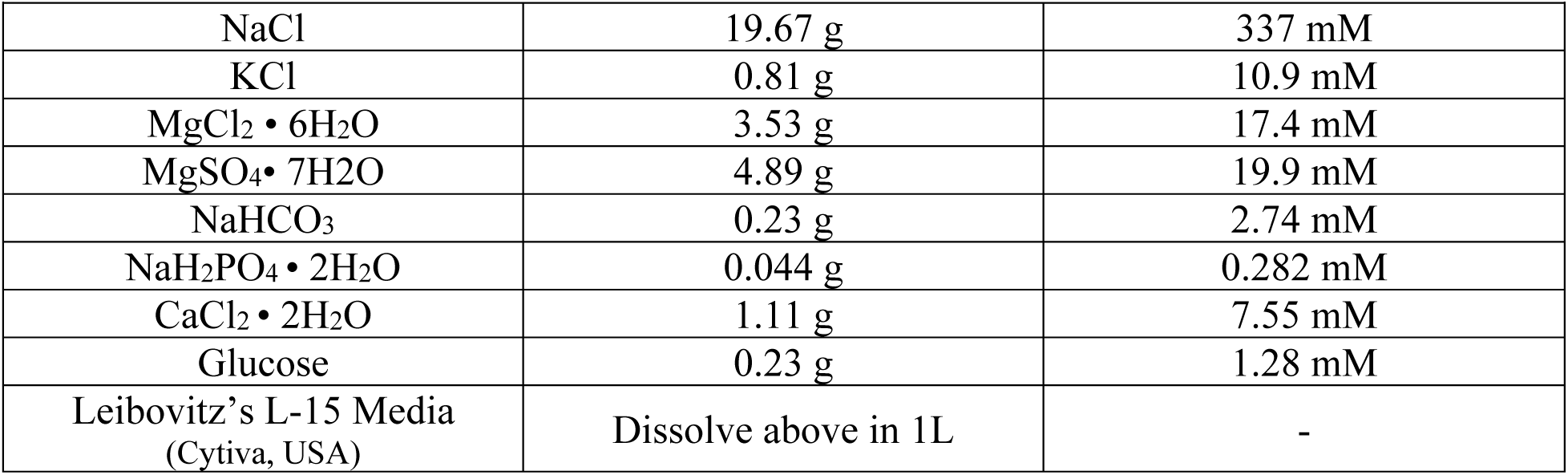
Base Media (1 Liter) ^15^

**Table 3:** Base Media w/Antibiotic & Complete Media (From Base Media in Table 2)15.

|  |  |
| --- | --- |
| **Base Media with Antibiotic** | 91.5% (v/v) Base Media |
|  | 8% (v/v) L-15 media |
|  | 0.5% (v/v) Penicillin-Streptomycin |
| **Complete Media** | 90.5% (v/v) Base Media |
|  | 8% (v/v) L-15 media |
|  | 0.5% (v/v) Penicillin-Streptomycin (Pen-Strep)<br>(10,000 U/mL) (Cytiva, USA) |
|  | 1% (v/v) Insulin-Transferrin-Selenium (ITS-G)<br>(Gibco, Thermo Fisher Scientific, USA) |
|  | 10 ng/mL of the fibroblast growth factor (FGF)<br>(Gibco, Thermo Fisher Scientific, USA) |
|  | 100 ng/mL of the epidermal growth factor (EGF)<br>(Gibco, Thermo Fisher Scientific, USA) |

### Hoechst and Phalloidin Cell Staining

Fresh working stocks of phalloidin (Sigma-Aldrich, USA) were prepared the day of the experiment by diluting 1 µL of the 1000x phalloidin conjugate stock solution in 1 mL of PBS-/-containing 1% BSA, and mixing thoroughly by pipetting. Cells were fixed in 4% paraformaldehyde (PFA) (Sigma-Aldrich, USA) in PBS for 15–20 minutes, followed by three washes with PBS-/-. Permeabilization was performed by incubating cells in 0.1% Triton X-100 in PBS-/- for 5 minutes, followed by three additional PBS-/- washes. A total of 100 µL of 1x phalloidin conjugate working solution was added to each sample, and cells were incubated at room temperature for 90 minutes. Following incubation, the solution was aspirated, and cells were stained with Hoechst 33342 (Sigma-Aldrich, USA) for 15-20 minutes. Samples were then washed three times with PBS-/-. Imaging for phalloidin and Hoechst staining was performed using excitation at 355 nm (Hoechst) and 488 nm (phalloidin) on a Zeiss Axio Observer 7 (Carl Zeiss, Germany) widefield microscope. All images were processed using FIJI (v2.9.0/1.53t) according to standard practices^30^.

### Flow Cytometry

Cells were washed in PBS-/- and centrifuged at 700 rcf. The resulting pellet was resuspended in PBS-/- and fixed in 70% cold ethanol (4 °C) for 30 min. Fixed cells were stored at 4 °C for 24 hours. Fixed cells were centrifuged at 700 rcf for 5 min at 4 °C, resuspended in cold PBS, and washed twice more under the same conditions. Prior to staining, cells were treated with RNase A (Sigma-Aldrich, USA) to remove RNA contamination and ensure accurate DNA content analysis. The fixed cells were incubated in the Propidium Iodide (Sigma-Aldrich, USA) staining solution (Table 4) at 37°C for 15 min, then transferred to ice or stored at 4 °C protected from light. Prior to analysis, samples were passed through flow cytometry tubes equipped with cell-strainer caps with a mesh size of 40 µm to remove debris and cell aggregates and analyzed the same day on a FACSCalibur Flow Cytometer (BD Biosciences, USA). Up to 20,000 events were collected on the FL2 channel (488 nm laser that detects at 575 nm) for each stained sample. A sequential gating strategy was applied in FlowJo (v10; BD Biosciences, USA), including forward scatter (FSC) and side scatter (SSC) to exclude debris, followed by doublet discrimination using FSC-A versus FSC-H to isolate single cells. Cell cycle analysis was performed on FlowJO (V10) (BD Biosciences, USA) using the Watson-pragmatic model to find the number of cells in G0/G1, S phase, and G2/M phase^31^. Flow cytometry experiments were performed using three biological replicates, each with three technical replicates.

**Table 4:**
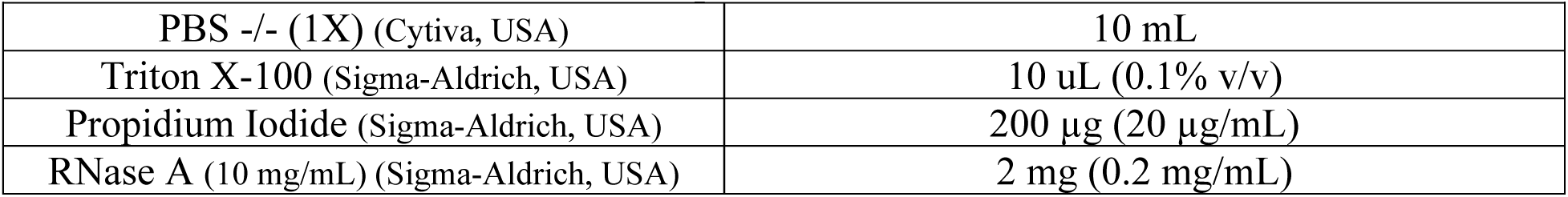
Propidium Iodide Solution.

## Acknowledgements

We thank the Marine Biological Laboratory’s Cephalopod Program team including Bret Grasse, Taylor Sakmar, and Miranda Vogt, for their extensive assistance with animal husbandry, euthanasia protocols, and the provision of octopus for the experiments. We also thank Dr. Yoonji Kim for meeting with our laboratory, sharing the protocol she developed in squid primary cultures to develop our work with octopus samples, and providing training in cephalopod optical lobe dissection. We would also like to thank the Life Science Editors Foundation’s JEDI award initiative and Dr. Stephen Matheson for assistance in development of a roadmap to this preprint.

## Author Contributions

J.V.P., S.A., and M.H. designed the study. J.V.P. and S.A. optimized octopus primary cell culture conditions. J.V.P., S.A., F. K. and P. S. performed octopus experiments and analysis, with guidance from M.H. J.V.P. wrote the initial draft of the manuscript. All authors contributed to the edits of the final text and the figures.

## Data Availability Statement

All data generated and analyzed during this study are included in this published article.

---

**Supplementary Figure 1:**
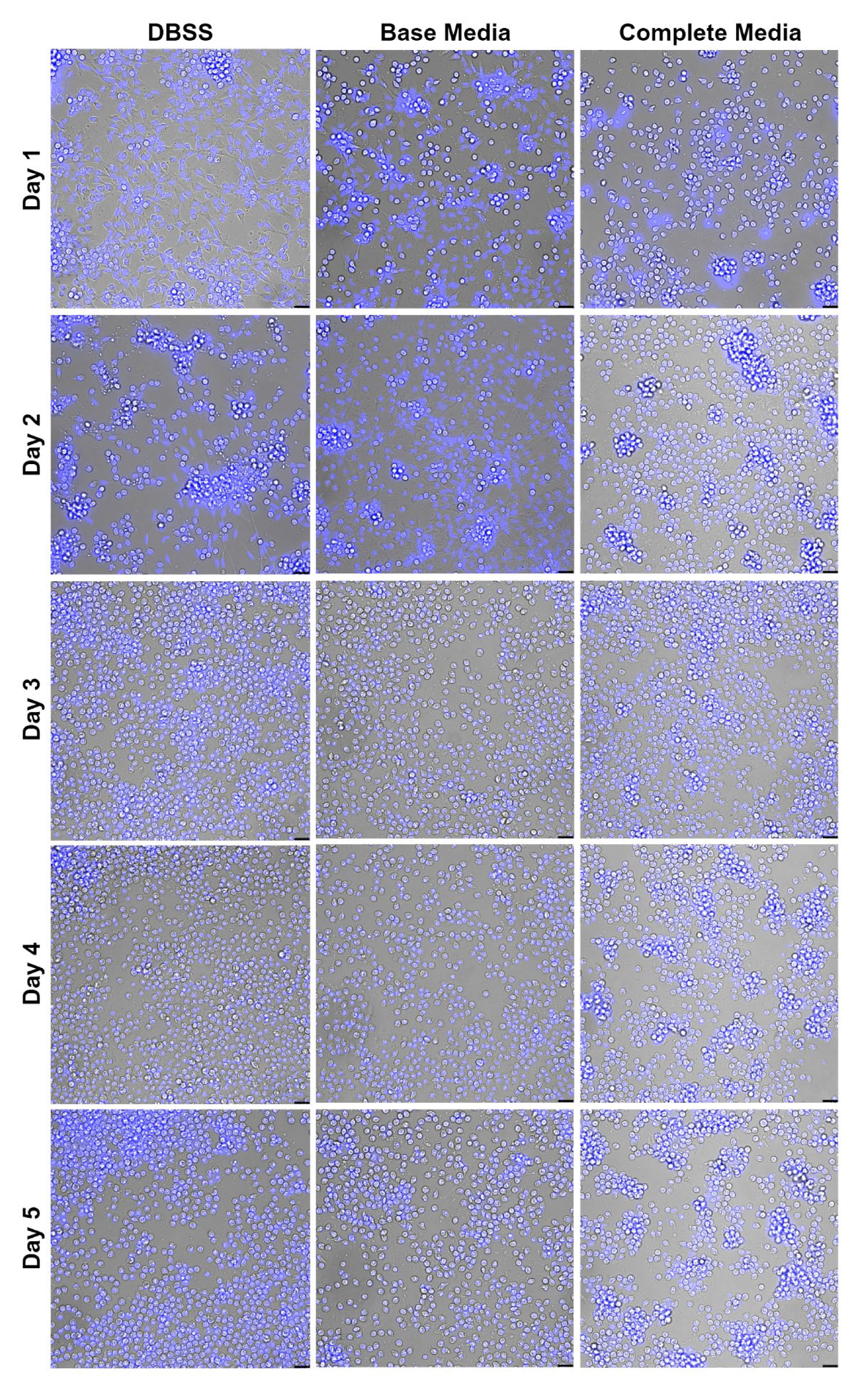
Time-course of primary cell cultures derived from *O. bimaculoides* brain optic lobe tissue under different media conditions. Representative brightfield images of dissociated cells cultured in DBSS, Base Media, or Complete Media over five days (Day 1–Day 5). Cells show progressive dissociation, survival, and aggregation dynamics depending on media composition. In DBSS, cells exhibit limited long-term viability with increasing debris over time. Base Media supports partial cell survival and moderate clustering. Complete Media promotes higher cell density, viability, and formation of multicellular aggregates across the culture period. Images are shown at consistent magnification; scale bars = 20 µm.

**Supplementary Figure 2:**
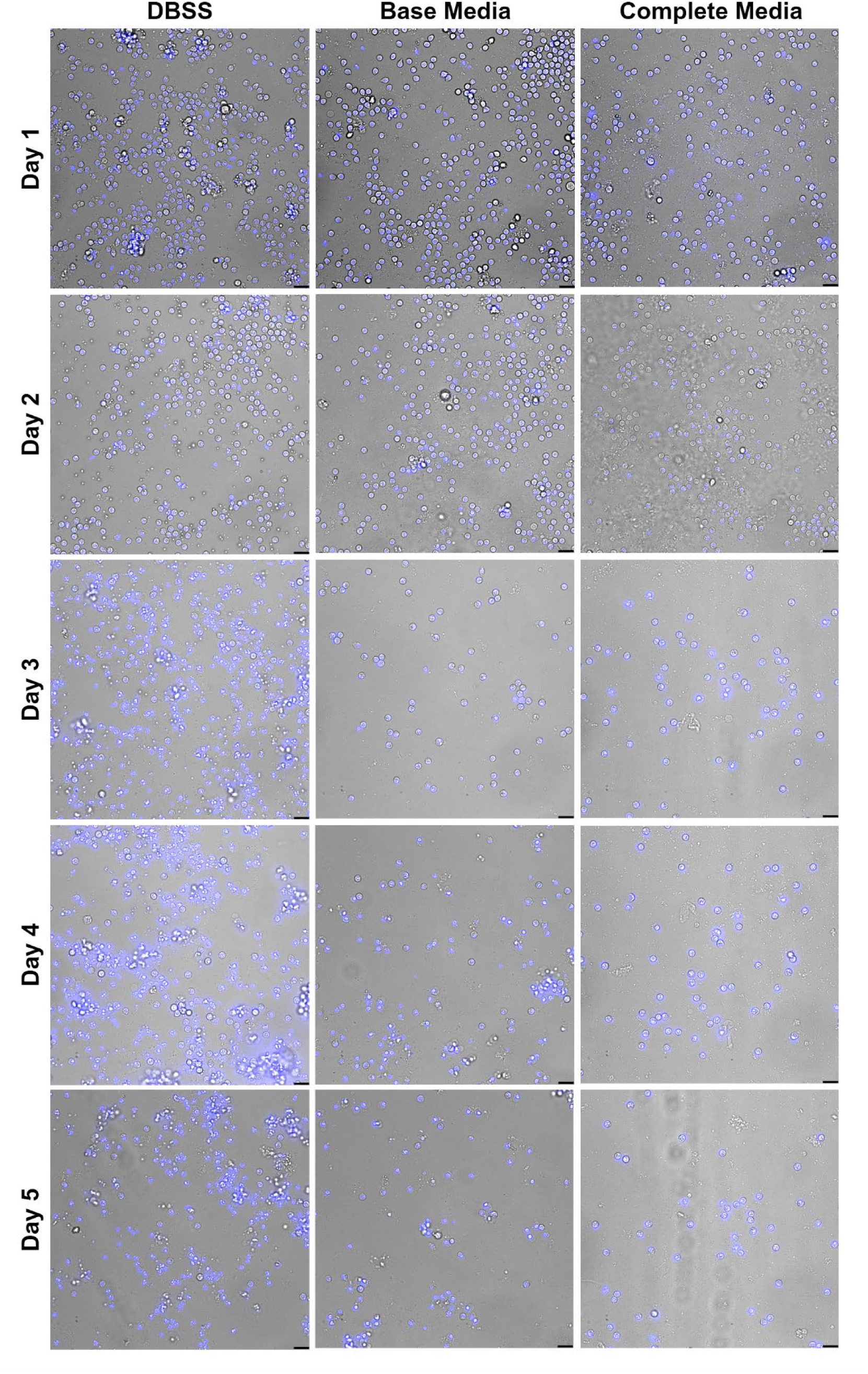
Time-course of primary cell cultures derived from *O. bimaculoides* venom posterior salivary gland tissue under different media conditions. Representative brightfield images of dissociated PSG-derived cells cultured in DBSS, Base Media, or Complete Media over five days (Day 1–Day 5). Across all conditions, cells progressively decrease in density over time, indicating reduced survival compared to brain-derived cultures. DBSS shows rapid decline with increased debris and loss of intact cells. Base Media supports limited short-term survival, while Complete Media modestly improves cell maintenance but does not sustain high cell density over multiple days. Images were acquired at consistent magnification; scale bar = 20 µm.

**Supplementary Figure 3:**
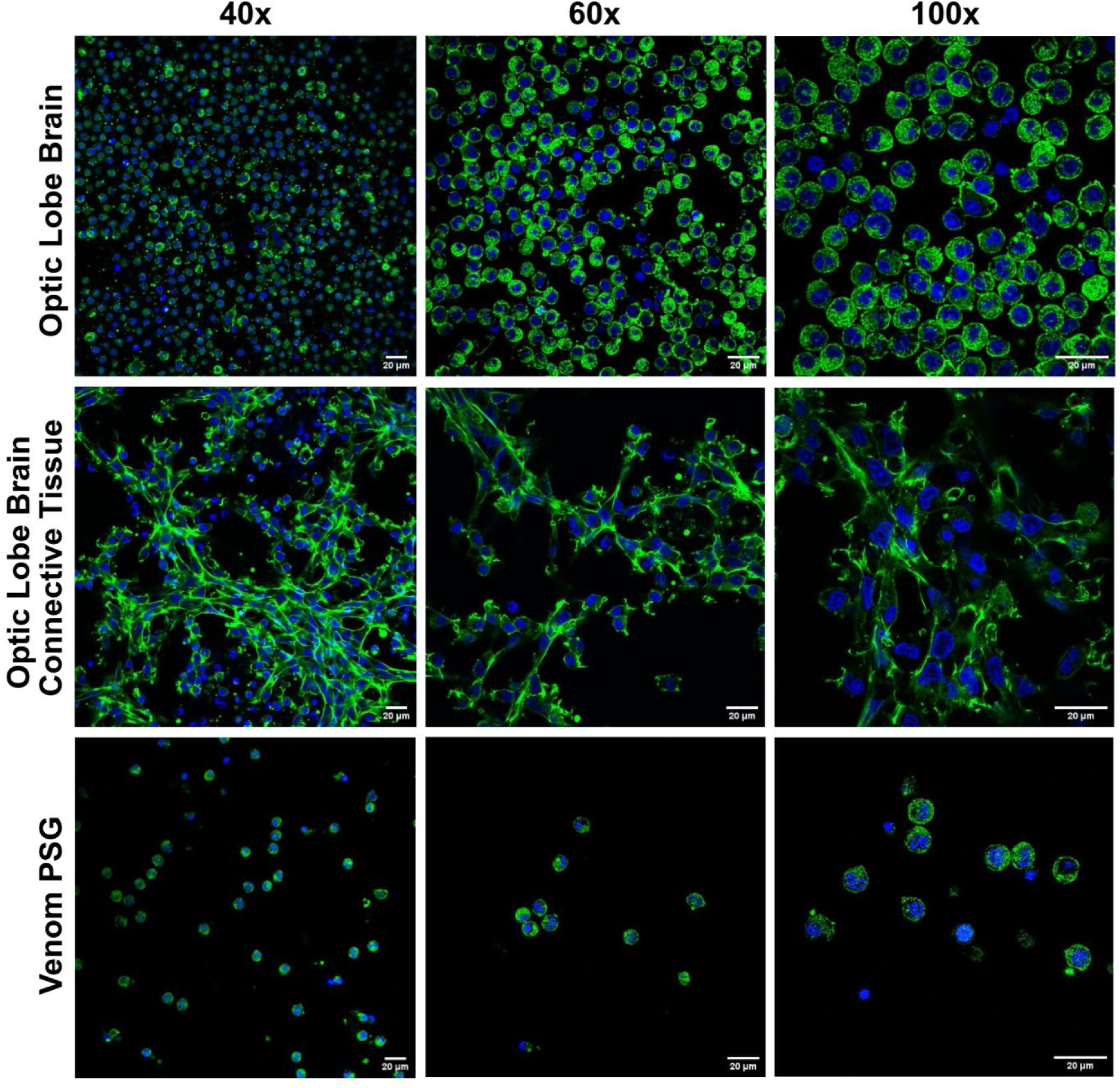
F-actin organization visualized using phalloidin and DAPI staining in dissociated brain optic lobe and venom PSG cells of *O. bimaculoides*. Representative fluorescence images of dissociated brain optic lobe, brain optic lobe connective tissue, and posterior salivary gland (PSG) cells imaged at 40x, 60x, and 100x magnification (left to right). Cells were stained with Hoechst (blue) to label nuclei and phalloidin (green) to visualize F-actin. Brain-derived cells appear predominantly rounded with cortical actin localization, whereas brain optic lobe connective tissue cells exhibit extended, network-like morphologies with prominent actin filaments. Scale bars, 20 µm.

